# On the Subspace Invariance of Population Responses

**DOI:** 10.1101/361568

**Authors:** Elaine Tring, Dario L. Ringach

## Abstract

The response of a neural population in cat primary visual cortex to the linear combination of two sinusoidal gratings (a plaid) can be well approximated by a weighted sum of the population responses to the individual gratings – a property we refer to as subspace invariance. We tested subspace invariance in mouse primary visual cortex by measuring the angle between the population response to a plaid and the plane spanned by the population responses to its individual components. We found robust violations of subspace invariance, represented by a median angular deviation of ~55 deg. The cause of this departure is a strong, negative correlation between the mean responses a neuron to the individual gratings and its response to the plaid. We suggest that an early nonlinearity may distort the power distribution of grating and plaid stimuli such that plaids have a prominent power component at ±45 deg off the fundamental orientations. We conclude that subspace invariance does not hold in mouse V1. This finding rules out a large class of possible models of population coding, including vector averaging and gain control.

## Introduction

How do population of neurons in primary visual cortex encode the visual input (1-7)? Are there elementary properties that would allow us to predict the response of a neural population to complex visual stimuli from the measurement of its responses to simple ones? Previous work measured the response of neural populations to gratings of different contrasts and the plaids that result from their superposition (8, 9). The population response to plaids were well described by a linear mixture of the responses to the individual component gratings. The mixing coefficients were explained by a contrast gain model (10). The model provided a satisfactory account for the averaging behavior observed when gratings had similar contrasts and the gradual transition to a winner-take-all regime as the contrasts became increasingly different (8).

The contrast gain model applied to population responses is an example of a system that satisfies *subspace invariance.* Given two stimuli, *A* and *B*, and their corresponding population responses, *r*(*A*) and *r*(*B*), we say the population exhibits subspace invariance if the response to a linear combination *C* of *A* and *B* is a linear combination of *r*(*A*) and *r*(*B*). In other words, given *C* = *αA* + *βB*, subspace invariance is satisfied if *r*(*C*) ∈ span(*r*(*A*), *r*(*B*)). We can test for subspace invariance by measuring the angle *φ* between *r*(*C*) and its best approximation lying in the plane *S* = span(*r*(*A*), *r*(*B*)) (**Fig 1**). Of course, the optimal approximation in the mean-squared sense is the projection of *r*(*C*) onto *S*, which we denote by *r͂*(*C*) = *P_s_r*(*C*). Here, we used this approach to test the subspace invariance of neural population responses in mouse primary visual cortex.

**Fig 1.**
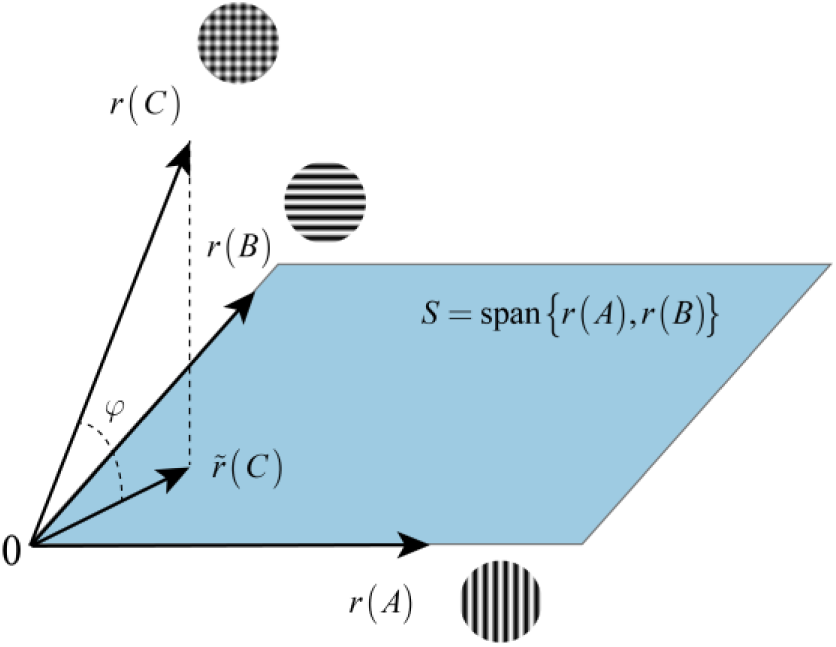
Subspace invariance. A population response is said to satisfy subspace invariance if the response to a linear combination of stimuli lies in the space spanned by the population responses to the individual stimuli. Departures from subspace invariance can be measured by the deviation angle *φ*, between the actual population response to a linear combination and by the best approximation resulting by a linear combination of the responses to each of the components.

## Methods

### Animals

All procedures were approved by UCLA’s Office of Animal Research Oversight (the Institutional Animal Care and Use Committee) and were in accord with guidelines set by the US National Institutes of Health. A total of 9 mice, both male (2) and female (7), aged P35-56, were used in this study. We used TRE-GCaMP6s line G6s2 (Jackson Lab) which express where GCaMP6s is regulated by the tetracycline-responsive regulatory element (tetO). Mice were housed in groups of 2-3, in reversed light cycle. Animals were naïve subjects with no prior history of participation in research studies. We imaged 36 different populations to obtain the data discussed in this study.

### Surgery

Carprofen and buprenorphine analgesia were administered pre-operatively. Mice were then anesthetized with isoflurane (4-5% induction; 1.5-2% surgery). Core body temperature was maintained at 37.5C using a feedback heating system. Eyes were coated with a thin layer of ophthalmic ointment to prevent desiccation. Anesthetized mice were mounted in a stereotaxic apparatus. Blunt ear bars were placed in the external auditory meatus to immobilize the head. A portion of the scalp overlying the two hemispheres of the cortex (approximately 8mm by 6mm) was then removed to expose the underlying skull. After the skull is exposed, it was dried and covered by a thin layer of Vetbond. After the Vetbond dries (~15 min) it provides a stable and solid surface to affix the aluminum bracket with dental acrylic. The bracket is then affixed to the skull and the margins sealed with Vetbond and dental acrylic to prevent infections.

### Imaging

We began imaging sessions ~4-5 days after surgery. We used a resonant, two-photon microscope (Neurolabware, Los Angeles, CA) controlled by Scanbox acquisition software (Scanbox, Los Angeles, CA). The light source was a Coherent Chameleon Ultra II laser (Coherent Inc, Santa Clara, CA) running at 920nm. The objective was an x16 water immersion lens (Nikon, 0.8NA, 3mm working distance). The microscope frame rate was 15.6Hz (512 lines with a resonant mirror at 8kHz). Eye movements and pupil size were recorded via a Dalsa Genie M1280 camera (Teledyne Dalsa, Ontario, Canada) fitted with a 740 nm long-pass filter that looked at the eye indirectly through the reflection of an infrared-reflecting glass. Images were captured at an average depth of 220 μm. Locomotion was monitored via a quadrature encoder coupled to the shaft of the wheel.

### Visual stimulation

In all experiments we used a BenQ XL2720Z screen which measured 60 cm by 34 cm and was viewed at 20 cm distance, subtending 112 × 80 degrees of visual angle. The screen was calibrated using a Spectrascan PR-655 spectro-radiometer (Jadak, Syracuse, NY), and the result used to generate the appropriate gamma corrections for the red, green and blue components via an nVidia Quadro K4000 graphics card. The contrast of individual sinusoidal gratings was 80%. The plaid was the sum of two gratings with a contrast of **80**%/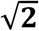
**56.6**%, to ensure the same effective contrast as the individual gratings. The spatial frequencies of the components was always the same, and ranged from 0.03 to 0.06 cpd. Visual stimuli were generated in real-time by a Processing sketch using OpenGL shaders (see http://processing.org) Transistor-transistor logic (TTL) signals generated by the stimulus computer were sampled by the microscope and time-stamped with the frame and line number that being scanned at that time. The time-stamps provided the synchronization between visual stimulation and imaging data. The center of the monitor was positioned with the center of the receptive field population for the eye contralateral to the cortical hemisphere under consideration. The approximate locations of the receptive fields of the population were estimated by an automated process where localized, flickering checkerboards patches, appeared at randomized locations within the screen. This experiment was run at the beginning of each imaging session to ensure the centering of receptive fields on the monitor. We imaged the monocular region of V1 in the left hemisphere. The receptive fields of neurons were centered around 20 to 35 deg in azimuth and 0 to 20 deg in elevation on the right visual hemifield.

### Image processing

The image processing pipeline was the same as described in detail elsewhere (11). Briefly, calcium images were aligned to correct for motion artifacts. We then used a Matlab graphical user interface (GUI) tool developed in our laboratory to define regions of interest corresponding to putative cell bodies manually. Following segmentation, we extracted signals by computing the mean of the calcium fluorescence within each region of interest and discounting the signals from the nearby neuropil. Spikes were then estimated via deconvolution (12).

### Data selection

The mean temporal response of each cell for each condition was computed (**Fig 2C**). We considered a cell to have generated a significant response to a given condition (one of the two gratings or the plaid) if the peak relative change in fluorescence over was larger than eight. Our findings are robust to changes in this criterion. We defined cell populations as the set of all cells which generated a significant response for at least one condition, all other neurons were discarded.

**Fig 2.**
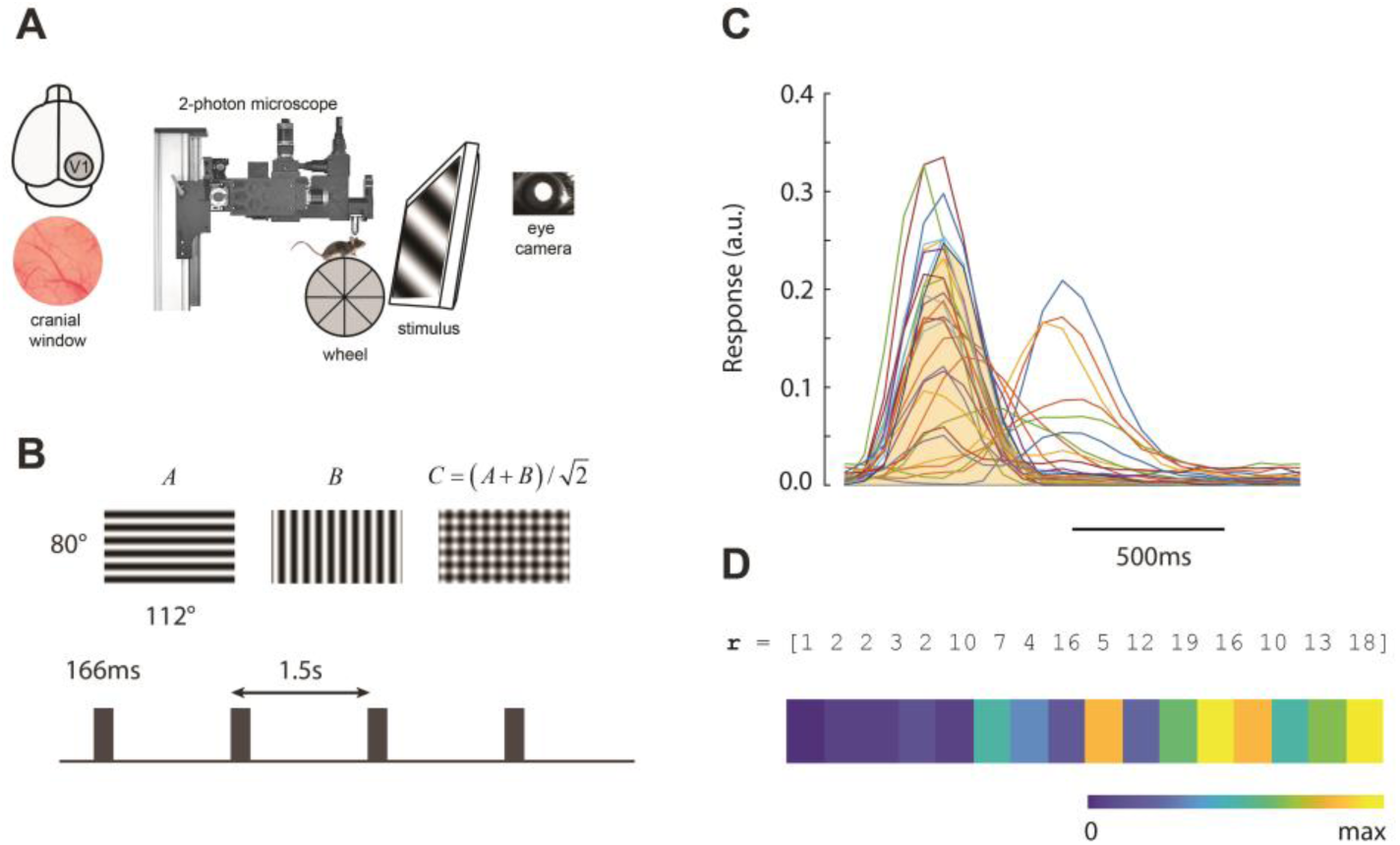
Experimental setup. (A) A cranial window is implanted on top of V1. Awake mice are imaged during the presentation of visual stimuli. Recordings of eye movements and locomotion accompany the imaging data. (B) Visual stimuli consist of two orthogonal sinusoidal gratings and a plaid resulting from their combination. Patterns are flashed briefly in random order. (C) Cortical responses return to baseline at the end of the 1.5 trial. The activity of each cell is represented by its integrated activity (shaded area). (D) Population vector responses are represented as a “barcode”.

### Code and Data availability

All analyses were conducted in Matlab (Mathworks, Natick, MA). Code and data will be available on Figshare.

## Results

We measure the activity of neural populations in primary visual cortex of awake, adult mice of both genders, expressing a calcium indicator (GCaMP6s) during exposure to computer-controlled visual stimulation (**Fig 2A**). Mice are head restrained, but otherwise free to walk, rest or groom on a 3D printed wheel. The stimulus sequence contained three different patterns: two sinusoidal gratings with the same spatial frequency and orthogonal orientations (patterns *A* and *B*), and a plaid resulting from their combination and the same effective contrast (pattern *C* = (*A* + **B**)
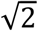)
(**Fig 2B**). In each trial, one of the patterns was selected at random with equal probability and flashed for 166ms, with a stimulus-onset asynchrony of 1.5s (**Fig 2B**). The responses of individual cells to such brief stimuli return to baseline by the end of the trial (**Fig 2C**). In the present study, we define the response of a cell as its integrated activity over the entire trial (**Fig 2C**, shaded area). We run 30 min long sessions for a total of 1200 trials (400 average presentations per pattern). We computed the average response for each pattern across trials, *r*(*A*), *r*(*B*), and *r*(*C*). The dimension of the responses equals the number of cells in the population after cell selection (see **Methods**). We find it convenient to represent the mean responses as a “barcode” where the response of each cell is mapped to a normalized colormap (**Fig 2C**).

To make it easier to visually interpret a population barcode we ordered the cells according to their relative preference for the two grating stimuli. Denote by *r_i_*(*A*) and *r_i_*(*B*) the mean response of the *i* – *th* cell in the population to patterns *A* and *B* respectively. Then, barcodes display the population response when cells are ordered increasingly by *r_i_*(*A*) – *r_i_*(*B*). In this representation, cells with preference for *A* over *B* will be on the right hand side of the barcode, while cells with a preference for *B* over *A* will be on the left.

For each experimental session we visualized the results by displaying the barcodes representing the mean population responses to each grating *r*(*A*) and *r*(*B*), the mean population response to the plaid, *r*(*C*), and the best approximation to *r*(*C*) attainable by a linear combination of *r*(*A*) and *r*(*B*), which we denoted by *r̃*(*C*) (**Fig 3A**). Due to our choice of cell ordering, the barcode for *r*(*A*) shows the most active cells are on the right, while the most active cells in the barcode for *r*(*B*) are on the left. Cells that are active during the presentation of a grating, however, appear not to be respond strongly to the plaid, as both ends of the barcode for the plaid are dominated by blue hues, corresponding to low firing rates. In contrast, cells responsive to the plaid are positoned somewhere in the middle of the barcode for *r*(*C*). These cells do not seem to respond to the individual components. Moreover, simple visual inspection reveals that the population response to a plaid is very different from its best approximation by a linear combination of the population responses to the components, violating the prediction of subspace invariance (**Fig 3A**).

**Fig 3.**
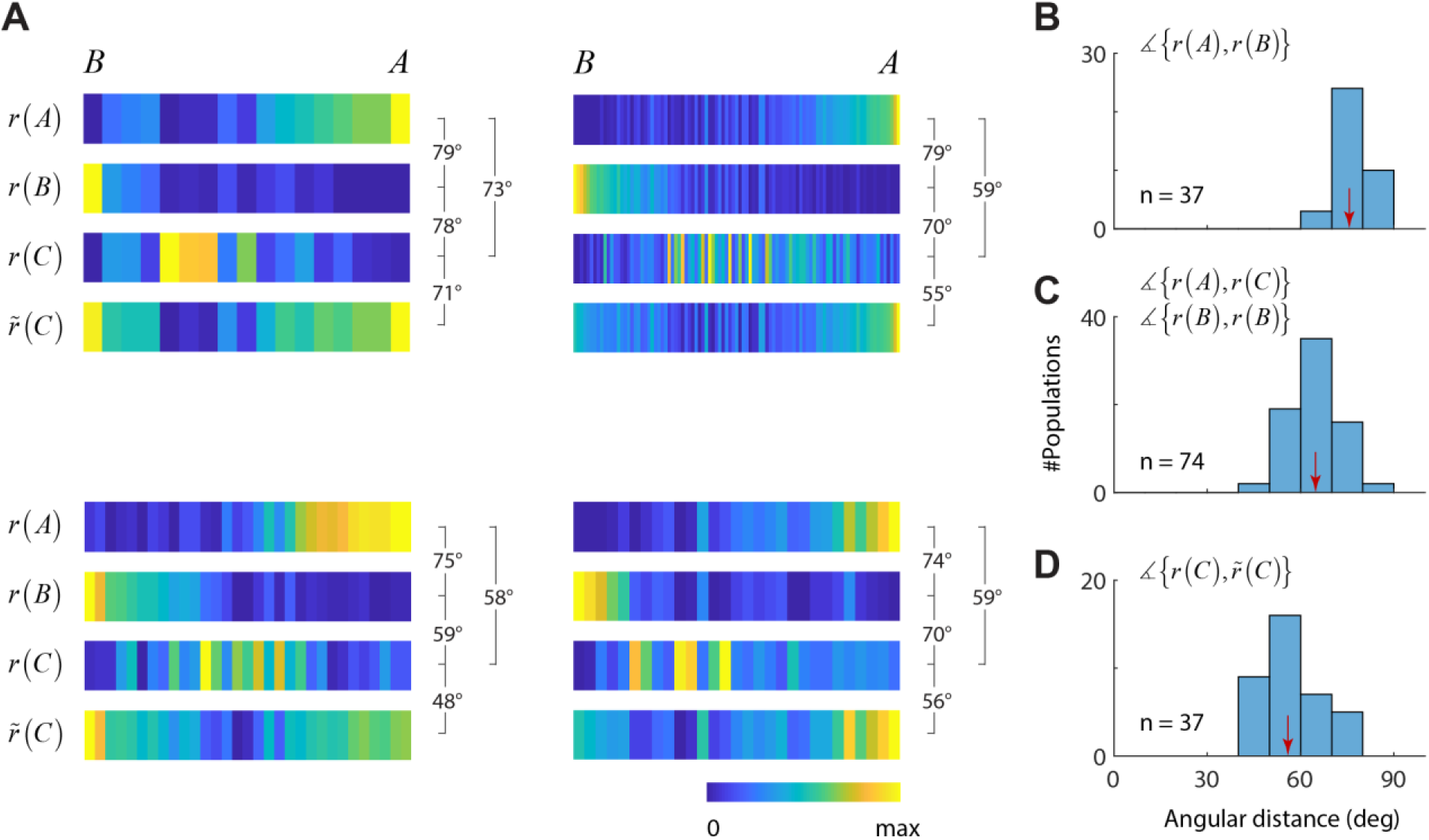
Testing subspace invariance in mouse V1. (**A**) Each panel shows the results from separate experiments. Within each panel, the barcodes display the mean population responses to the gratings, *r*(*A*) and *r*(*B*), the mean response to the plaid, *r*(*C*), and the best approximation to the population response to the plaid that lies in the plane spanned by *r*(*A*) and *r*(*B*), denoted by *r͂*(*C*). The brackets indicate the angular distance between the different population vectors. (**B**) Distribution of angular distance between mean population responses to the sinusoidal gratings. (**C**) Distribution of angular distance between mean population responses to the sinusoidal gratings and the plaid. (**D**) Distribution of angular distance between the response to the plaid and the best approximation in the plane spanned by the grating responses. (**B-D**) Red arrows indicate median angular distances in each case.

We quantified these observations by computing the angular distance between mean population vectors (**Fig 3B-D**). Across all experiments, the mean angular distance between the population responses to the gratings was large, with a median of 75.4 deg (**Fig 3B**). The median angular distance between the population response to the gratings and the plaid stimulus was 64.2 deg (**Fig 3C**). Importantly, the median distance between the population response to the plaid and its projection on the subspace spanned by the responses to gratings was 56.5 deg. The medians of the distributions were all statistically different at the *p* < 10^−4^ level.

A different way to express the deviations from subspace invariance is to calculate the relative error, defined as the norm of the difference between the actual plaid response and its projection, normalized by the norm of the plaid response, *ϵ* = ∥*r*(*C*) – *r̃*(*C*)∥/∥*r*(*C*)∥ (**Fig 4A**). The normalized error is relatively large, with a median of 0.83. These data are effectively the same as shown in **Fig 3D**, as one can easily see that *ϵ* = sin *φ*(**Fig 1**). These large deviations are not the result of weak population responses to the plaid, as the magnitude of the population responses to the gratings and the plaids were comparable (**Fig 2B**). In fact, plaids generted responses that were 1.15 times larger than the mean norm of the responses to the gratings.

**Fig 4.**
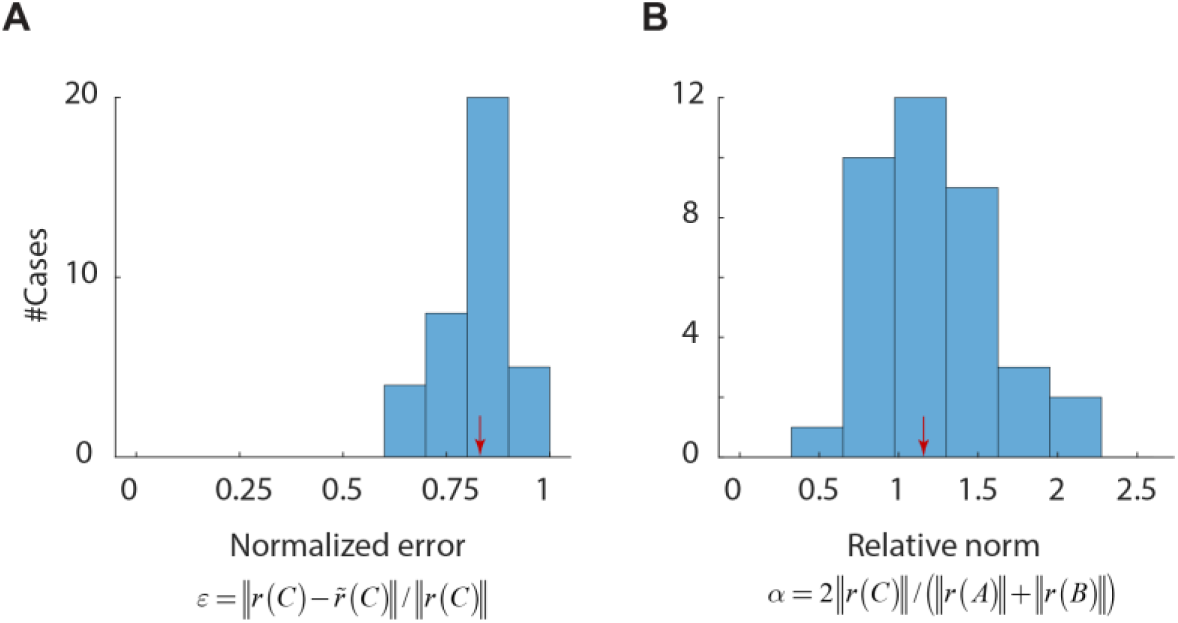
(**A**) Histogram of normalized error for all experiments in the dataset. (**B**) Relative magnitude of the plaid population responses with respect to those evoked by gratings.

To compute an “average barcode” across all our datasets we first normalized the width of the barcodes to one and considered the values of the barcode as samples of a function in that interval. We then interpolated these data at 32 equally spaced points and averaged the results for each stimulus class to obtain average barcodes and angular distances (**Fig 5A**). These composite barcodes highlight the difference in the population responses for each stimulus condition. It also accentuates one key feature of the data already noted in the individual cases (**Fig 3A**) – namely, cells in the population that respond to the plaids do not respond to the individual components, while cells that respond to the individual components do not respond to the plaid and viceversa. This is the fundamental reason why it is not possible to approximate the population response to the plaid as a linear combination of the responses to the gratings. This effect, which is so salient in the average data, is also present if we look at the responses of individual cells (**Fig 5B**). The scatterplot shows the average response of a cell to the plaid, *r_i_*(*C*), against its mean response to the gratings, (*r_i_*(*A*) + *r_i_*(*B*))/2. All individual cells recorded are included in this analysis. The responses of individual cells to plaids and gratings are negatively correlated (*n* = 1035, *r* = –0.62, *p* < 10^−100^).

**Fig 5.**
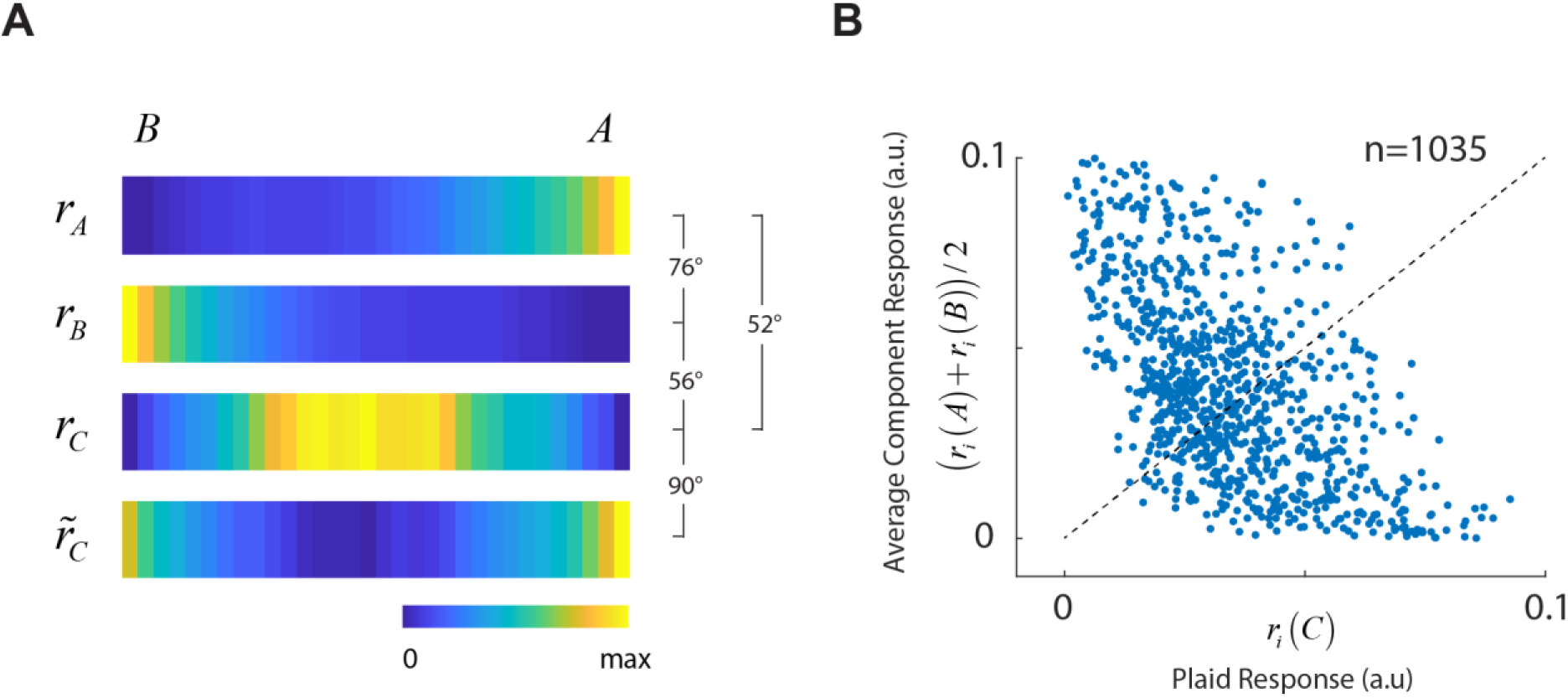
Average barcodes and behavior of single neurons. (**A**) Average barcodes across all our experiments. As before cells are ordered according to their preference between gratings *A* and *B*. The average barcodes highlight the differences between the population responses to gratings and plaids. The best approximation to the plaid population from linear combination to the average response to gratings is poor. (**B**) A scatterplot of plaid versus average responses to gratings for each individual cell shows a statistically significant, and strong anticorrelation. Dotted line represents the unity line.

What kind of mechanism could generate the negative correlation between component responses to sinuoidal gratings and plaids? One explanation is that there is a front-end nonlinearity that distorts the contrast of visual patterns. A strong nonlinearity will affect the distribution of power in the Fourier domain (**Fig 6**). While the spectra of pure sinusoidal gratings and plaids will consist of peaks at the fundamental frequencies (**Fig 6A**), a sigmoidal distorsion of the contrast will spread the spectral components to include higher harmonics. In the case of the gratings, the harmonics will have the same orientation as the fundamentals. However, in the case of the distorted plaid, the dominant power will be contained in harmonics at ±45 deg off the fundamental orientations (13). It is then possible for cells that respond to an appropiate range of orientation and spatial frequencies to respond to the plaid and not the grating, and viceversa. To test this idea we are currently collecting data that includes not only the mean responses to gratings and plaids, but also the full 2D tuning of neurons in the orientation and spatial frequency domain. A comparison of the tuning of neurons at different positions of the barcode should be able to tell us if the proposed explanation is likely or not. At present we do not have sufficient data to make a determination.

**Fig 6.**
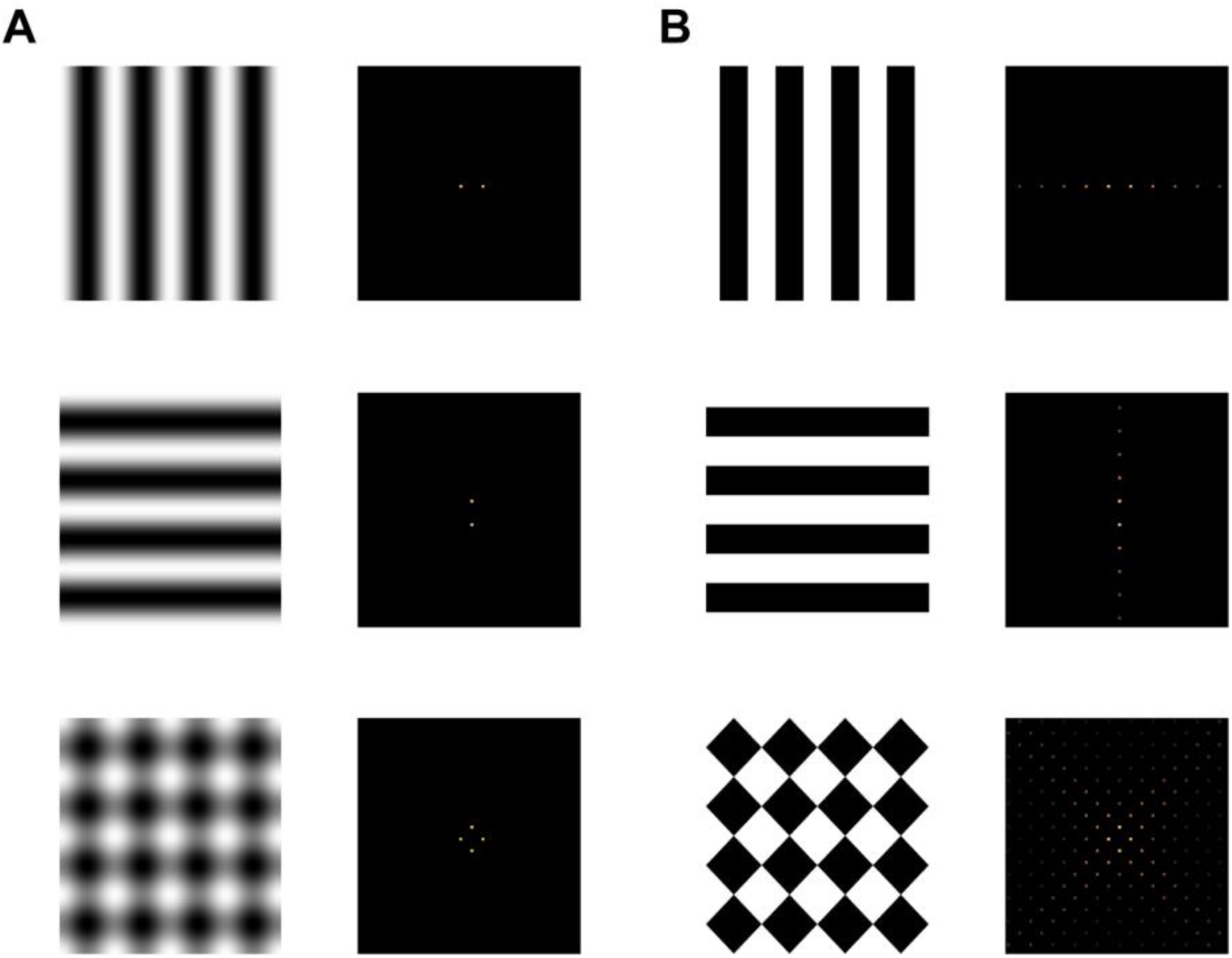
Early static nonlinearities generate higher harmonics at ±45 deg off the fundamental orientations of the grating components. (**A**) Undistorted sinusoidal gratings and the resulting plaid had a power distribution concentrated at the fundamental frequencies. (**B**) A sigmoidal distortion of the contrast spreads the power spectrum across higher harmonics. In the case of the gratings all harmonics have the same orientation as the grating, but in the case of the plaid, the power is concentrated on harmonics at ±45 deg off the fundamentals.

## Discussion

Understanding how populations of neurons respond to natural stimulation is a central problem in systems neuroscience (1, 7). Here we studied subspace invariance of population response, which is a weaker property than the homogeneity of linear systems, as it does not require strict superposition of the responses. In a linear system *r*(*αA* + *βB*) = *αr*(*A*) + *βr*(*B*), while subspace invariance only requires that *r*(*αA* + *βB*) ∈ span{*r*(*A*), *r*(*B*)}. An example of a non-linear model which satisfies subspace invariant is the normalization model (8), which when applied to population of neurons in the primary visual cortex of the cat, was able to explain rather well the responses to plaid stimuli composed of gratings of different contrast levels. The model accounts for the averaging behavior observed for gratings of equal contrasts and the transition into a winner-take-all regime as the contrasts increasingly diverge, making the identity of the stimulus with highest contrast determine the population response. A second study used intrinsic imaging in tree shrew primary visual cortex and reported averaging behavior of component with equal contrast matched the responses to plaids (9). Of course, averaging also satisfies subspace invariance.

In contrast to these past studies in cats and tree shrews, we find gross violations of subspace invariance in mouse primary visual cortex. The responses of neural populations to plaids could not be approximated by the linear combination of the population responses to its components, as evidenced by a large, median angular deviation of 56.5 deg and a relative error of 0.83 (**Fig 3D & 4A**). This result falsifies the hypothesis of subspace invariance and, as a special case, the gain control model as well. The central reason why subspace invariance fails is a strong, negative correlation between the average responses cells to the component gratings and to the plaid (**Fig 5B**). This means there were some cells that responded to the plaids that were unresponsive to the gratings.

Is it possible that subspace invariance holds for subthreshold responses but fails when analyzing extracellular responses? Let us assume that *r*(*A*) = *f*(*u*(*A*)), where *u*(*A*) is a subthreshold response to grating *A* and *f*(⋅) is a monotonically increasing, convex nonlinearity. We also adopt a similar notation for the other stimulus classes. Then, the response to a convex mixture of *A* and *B* satisfies *r*(*tA* + (1 – *t*)*B*) ≤ *t r*(*A*) + (1 – *t*) *r*(*B*) (Jensen’s inequality). For plaids constructed by averaging of two grating stimuli, we have *t* = 1/2 and *r*((*A* + *B*)/2)≤ (*r*(*A*) + *r*(*B*))/2. However, it is clear there are many cells for which this inequality if violated (**Fig 5B**). In particular, cells showing responses near zero to the gratings, should see their responses to the plaid bounded from above by very small number. Instead, these are the cells that show the maximal response to the plaid. Thus, a simple output nonlinearity is an unlikely explanation for the observed phenomenon. (Although our plaid stimuli are not strictly convex mixtures of the component gratings, as the norm of the stimulus constant, it is doubtful our findings would be any different had we chosen to average them instead.) Instead, we suggest that an early nonlinearity may distort the power distribution of grating and plaid stimuli such that plaids have a prominent power component at ±45 deg off the fundamental orientations. To assertain the validity of such explanation it would be necessary to measure the joint orientation of neurons for orinetation and spatial frequency, which is one of our next steps in this research project.

There are other differences in the stimuli and methods used across the studies. We only consider cells which generated a significant response to at least one class of stimuli. Thus, our barcodes, do not contain cells that do not respond to any of these stimuli. In contrast, the study by Busse and collaborators measured from cells which responded also to gratings at intermediate orientations than the ones corresponding to the gratings (8). Their population responses were defined first by arranging cells according to their preferred orientation and then averaging their responses in a discrete number of orientation bins. Assume the neurons that respond to plaids, but not gratings, have preferences for intermediate orientations when probed with gratings. One could also imagine that these cells represent a minority among other cells with similar orientation preferences. Then, this past analysis may have washed out the activities of cells responsive to plaids by averaging them with other non-responsive cells which have a similar preference for orientation. Another methodological difference is that cat recording occurred in a population of cells recorded with a 10×10 microelectrode array with a grid spacing of 400*μ*m. Such a population contains cells with pairwise distances much larger than the one in our two-photon data. It is possible that interactions between the members of the population that give rise to the negative correlations observed here are due to local network mechanisms not detectable when sampling from cell pairs which are far apart in the cortex.

Another possibility, of course, is that the results reflect true species differences. Although in our experiments stimuli were flashed gratings and plaids, we note that when probed with moving stimuli the proportion of component versus pattern cells in mouse appears different than those cats and monkeys (14-16) (but see ref (17) for a different outcome). According to one study (14), the fraction of component cells in mouse V1 is only 17%, compared to 84% in higher mammals, while the fraction of pattern cells is about 10%, compared to <1.3% in cats and monkeys. It would be of interest to see if there is any relationship between the responses of neurons to flashed gratings/plaids and the classic definition of component and pattern cells defined by moving stimuli (16, 18).

Altogether, we conclude the population responses in mouse V1 to the weighted sum of stimuli are not readily explained by a linear combination of their responses to the individual components. More complex models of population coding need to be developed. To this end, a deeper understanding between the relationship of cells that respond to plaids and their full tuning in the orientation and spatial frequency domain, and their categorization into component and pattern cells in response to moving stimuli, is needed. Finally, imaging the activity of LGN afferents should reveal if some of the properties observed is inherited from the input (19-22).

## References

1. Georgopoulos AP, Schwartz AB, Kettner RE. Neuronal population coding of movement direction. Science. 1986;233(4771):1416–9. PubMed PMID: 3749885.

2. Sompolinsky H, Yoon H, Kang K, Shamir M. Population coding in neuronal systems with correlated noise. Phys Rev E Stat Nonlin Soft Matter Phys. 2001;64(5 Pt 1):051904. PubMed PMID: 11735965.

3. Singh G, Memoli F, Ishkhanov T, Sapiro G, Carlsson G, Ringach DL. Topological analysis of population activity in visual cortex. J Vision. 2008;8(8). PubMed PMID: W0S:000258708900011.

4. Graf AB, Kohn A, Jazayeri M, Movshon JA. Decoding the activity of neuronal populations in macaque primary visual cortex. Nat Neurosci. 2011;14(2):239–45. doi:10.1038/nn.2733. PubMed PMID: 21217762; PMCID: PMC3081541.

5. Miller JE, Ayzenshtat I, Carrillo-Reid L, Yuste R. Visual stimuli recruit intrinsically generated cortical ensembles. Proc Natl Acad Sci U S A. 2014;111(38):E4053–61. doi:10.1073/pnas.1406077111. PubMed PMID: 25201983; PMCID: PMC4183303.

6. Lin IC, Okun M, Carandini M, Harris KD. The Nature of Shared Cortical Variability. Neuron. 2015;87(3):644–56. doi:10.1016/j.neuron.2015.06.035. PubMed PMID: 26212710; PMCID: PMC4534383.

7. Averbeck BB, Latham PE, Pouget A. Neural correlations, population coding and computation. Nat Rev Neurosci. 2006;7(5):358–66. doi:10.1038/nrn1888. PubMed PMID: 16760916.

8. Busse L, Wade AR, Carandini M. Representation of concurrent stimuli by population activity in visual cortex. Neuron. 2009;64(6):931–42. doi:10.1016/j.neuron.2009.11.004. PubMed PMID: 20064398; PMCID: PMC2807406.

9. MacEvoy SP, Tucker TR, Fitzpatrick D. A precise form of divisive suppression supports population coding in the primary visual cortex. Nat Neurosci. 2009;12(5):637–45. doi:10.1038/nn.2310. PubMed PMID: 19396165; PMCID: PMC2875123.

10. Heeger DJ. Normalization of cell responses in cat striate cortex. Vis Neurosci. 1992;9(2):181–97. PubMed PMID: 1504027.

11. Ringach DL, Mineault PJ, Tring E, Olivas ND, Garcia-Junco-Clemente P, Trachtenberg JT. Spatial clustering of tuning in mouse primary visual cortex. Nat Commun. 2016;7:12270. doi:10.1038/ncomms12270. PubMed PMID: 27481398; PMCID: PMC4974656.

12. Berens P. Community-based benchmarking improves spike inference from two-photon calcium imaging data. BioRxiv. 2017:doi: https://doi.org/10.1101/177956. doi: https://doi.org/10.1101/177956.

13. De Valois KK, De Valois RL, Yund EW. Responses of striate cortex cells to grating and checkerboard patterns. J Physiol. 1979;291:483–505. PubMed PMID: 113531; PMCID: PMC1280915.

14. Palagina G, Meyer JF, Smirnakis SM. Complex Visual Motion Representation in Mouse Area V1. J Neurosci. 2017;37(1):164–83. doi:10.1523/JNEUR0SCI.0997-16.2017. PubMed PMID: 28053039; PMCID:PMC5214628.

15. Muir DR, Roth MM, Helmchen F, Kampa BM. Model-based analysis of pattern motion processing in mouse primary visual cortex. Front Neural Circuits. 2015;9:38. doi:10.3389/fncir.2015.00038. PubMed PMID: 26300738; PMCID: PMC4525018.

16. Movshon JA, Newsome WT. Visual response properties of striate cortical neurons projecting to area MT in macaque monkeys. J Neurosci. 1996;16(23):7733–41. PubMed PMID: 8922429.

17. Juavinett AL, Callaway EM. Pattern and Component Motion Responses in Mouse Visual Cortical Areas. Current Biology. 2015;25(13):1759–64. doi:10.1016/j.cub.2015.05.028. PubMed PMID: WOS:000357360800029.

18. Adelson EH, Movshon JA. Phenomenal coherence of moving visual patterns. Nature. 1982;300(5892):523–5. PubMed PMID: 7144903.

19. Piscopo DM, El-Danaf RN, Huberman AD, Niell CM. Diverse visual features encoded in mouse lateral geniculate nucleus. The Journal of neuroscience. 2013;33(11):4642–56.

20. Zhao X, Chen H, Liu X, Cang J. Orientation-selective responses in the mouse lateral geniculate nucleus. J Neurosci. 2013;33(31):12751–63. doi:10.1523/JNEUROSCI.0095-13.2013. PubMed PMID: 23904611; PMCID: PMC3728687.

21. Kaye AP, Marshel JH, Nauhaus I, Callaway EM. Direction selectivity in the mouse LGN revealed by in vivo two-photon calcium imaging. Society for Neuroscience Abstract Viewer and Itinerary Planner. 2011;41. PubMed PMID: BCI:BCI201200082884.

22. Tang J, Ardila Jimenez SC, Chakraborty S, Schultz SR. Visual Receptive Field Properties of Neurons in the Mouse Lateral Geniculate Nucleus. PLoS One. 2016;11(1):e0146017. doi:10.1371/journal.pone.0146017. PubMed PMID: 26741374; PMCID: PMC4712148.

